# Creatine transporter (Slc6a8) knockout mice show an inattentive-like phenotype in the 5-choice serial reaction time test

**DOI:** 10.1101/2022.06.02.494536

**Authors:** Matthew R. Skelton, Rosalyn Liou, Marla K. Perna

## Abstract

Disorders of creatine (Cr) synthesis and transport cause moderate to severe intellectual disability, epilepsy, and a lack of speech development. Mutations of the X-linked Cr transporter (CrT; *SLC6A8*) gene are the most frequent cause of Cr deficiency and one of the leading causes of X-linked intellectual disability. There are no treatments for CrT deficiency (CTD) and there are many unanswered questions related to this disorder. Rodent models of CTD have deficits in spatial learning and memory, object recognition memory, fear conditioning, and working memory, making them high-fidelity models of CTD. While these cognitive deficits provide important information related to CTD, they lack some translational relevance and do not address important aspects of executive function like attention and impulsivity. To address this gap in knowledge, we tested brain specific *Slc6a8* knockout (bKO) mice in the 5-choice serial reaction time test (5CSRTT), a correlate of the continuous performance task in humans. Following 5CSRTT training, mice were then tested for 3 sessions using trials with a variable stimulus duration followed by 3 sessions using a fixed stimulus duration and variable intertrial interval. During both the testing phases the bKO mice had reduced accuracy along with increased omissions and correct latencies compared with controls. There were no increases in premature responses during the vITI, suggesting that these mice do not have an impulsive phenotype. The results of this study expand the known phenotype of *Slc6a8* deficient mice and add a translationally relevant behavioral output to test potential therapies.

**Synopsis:** This study shows that a mouse model of human creatine transporter deficiency, a devastating human condition, has an attention-deficit disorder-like phenotype without an increase in impulsivity.

## Introduction

Creatine (Cr) plays an essential role in maintaining ATP levels in cells with a high-energy demand^1-3^. The highest concentrations of Cr are observed in the skeletal muscle, heart, brain, and kidney^4^. Circulating Cr is taken into tissues using via the Cr transporter (CrT), which is encoded by the *SLC6A8* gene^5,6^. In cells, phospho-Cr (PCr) is formed when a phosphate group from ATP is donated to Cr via Cr kinase (CK) enzymes^7^. When required, PCr can donate this phosphate group to ADP via the CK mediated reverse reaction. This Cr-PCr shuttle acts as a spatial and temporal ATP buffer, providing a greater reserve capacity than the production of ATP alone^7^.

Cerebral Cr deficiency syndromes (CCDS) are a collection of disorders caused by the lack of Cr synthesis or transport^8^. They are characterized by severe to moderate intellectual disability (ID), epilepsy, autistic-like features, and aphasia^9,10^. The loss of the CrT, encoded by the *SLC6A8* gene, is the most prevalent CCDS. *SLC6A8* is located on the X-chromosome and CrT deficiency (CTD) shows an X-linked pattern of inheritance, with males being more affected than female carriers. Many CTD patients are also diagnosed with attention deficit hyperactivity disorder (ADHD). Prevalence studies suggest that CTD could account for 1-3% of all XLID ^11-16^, making it a leading known cause of ID. There is no treatment for CTD.

Several rodent models have been used to better understand the cognitive deficits related to CTD. Our lab developed the first model of CTD by removing exons 2-4 of the mouse *Slc6a8* gene^17^. These mice have robust deficits in spatial learning and memory, novel object recognition (NOR) memory, and contextual and conditioned fear memory^17-19^. A mouse lacking exons 5-7 of the *Slc6a8* gene was developed by Baroncelli et al had deficits in spatial learning and NOR deficits^20^. Working memory deficits were observed in these mice by measuring spontaneous alternation in the Y-maze. A rat model of CTD had reductions in spontaneous alternation; however, they did not show deficits in a spatial learning version of the Y-maze or in NOR memory^21^. Together, these data show that *Slc6a8* deficiency in rodents leads to cognitive deficits, suggesting they are high fidelity models of CTD. However, many of the learning tasks used do not have a direct human correlate, reducing their translational power. In addition, many of these behaviors focus on memory tasks and do not evaluate aspects of executive function. The purpose of this study was to evaluate attentional control in *Slc6a8*^*-/y*^ mice using a test with a high degree of translational value: the five-choice serial reaction time task (5C-SRTT)^22^. The 5C-SRTT is analogous to the human continuous performance task^23^. The 5C-SRTT measures aspects of executive function, including attention, impulsivity, and processing speed ^24-26^. Since many CTD patients are also diagnosed with ADHD, this test will provide further validation of our model and identify additional treatment benchmarks for this devastating disorder.

## Methods and Materials

### Subjects

The ubiquitous *Slc6a8*^*-/y*^ mice have significant body size reductions and there was a concern that they would not tolerate the food restriction protocol required for testing. Indeed, in a preliminary study the *Slc6a8*^*-/y*^ did not have the sufficient body mass and had to be rescued prior to the 10% weight loss required for testing. Therefore, we used the brain-specific *Slc6a8* knockout mice from Udobi et al that show learning deficits similar to those seen in the ubiquitous *Slc6a8*^*-/y*^ mice without the corresponding size reductions^18^. Female floxed *Slc6a8 (Slc6a8*^*Fl/+*^) mice were mated with male *Nestin-Cre* (Cre) mice as described by Udobi et al^18^ and the subjects of these pairings were tested. No more than 1 mouse per genotype was taken from a single litter. While every effort was made to use litter mates, not all genes were represented within a litter. All mice were on a C57BL/6J background. A total of 9 *Slc6a8*^*+/y*^ (WT), 10 Cre, 10 *Slc6a8*^*Fl/y*^ (Flox), and 14 *Slc6a8*^*Fl/y*^*::Nestin-Cre*^*+*^ (bKO) mice were used for this study. Prior to testing, mice were grouped housed in standard shoebox cages with *ad libitum* food and water. The housing room is kept on a 14:10 light:dark cycle with lights on at 0600 and the temperature is maintained at 21 ± 1º C.

### Food restriction

Two weeks prior to testing mice were moved from the standard housing room to the 5CSRTT housing room. The 5CSRTT housing room is maintained on a reverse light:dark cycle that matches the 14:10 duration of the holding room but the lights come on at 1600. Mice were transferred to the holding room in the afternoon as close to lights on as possible and housed individually for the duration of testing. Food restriction began one week after the transfer to the 5CSRTT housing room. Food was restricted on a gradual basis until mice reached 85-90% of their pre-testing weight. Testing was performed between 1000-1400 five days per week.

### Five Choice Serial Reaction Time Task

The 5C-SRTT was performed similar to the procedure outlined in Higgins and Silenieks ^27^. The 5C-SRTT apparatus is a 55.7 × 38.1 × 41 cm arena (Med Associates, St. Albans, VT) with 5 recessed square holes evenly distributed on the 55 cm testing wall and a square hole reward interface on the opposite reward wall. Each square hole is equipped with an LED light and an IR beam to detect entry. The apparatus is placed in a sound attenuated chamber with a fan to generate white noise. Within the reward interface a dipper is attached to a motorized arm that delivers ∼15 µL of the liquid reward (Strawberry Nesquik®). The units are controlled by the MedAssociates SmartCtrl package that allows 8 inputs and 16 outputs per chamber. Med-PC V Software Suite was used to program the behavioral tests executed by the chambers. The house light within the chamber was illuminated for testing.

In the first stage of training (Habituation 1), mice were familiarized to the chamber and the reward delivery by illumination of the reward light and raising the dipper every 10 s for 20 min. Head entries were recorded and the mice were advanced to the next stage (Habituation 2) following 2 consecutive days of >60 claimed rewards. During Habituation 2, mice initiated the testing session by claiming a free reward from the illuminated reward receptacle. After 4 s, all five lights on the training wall were illuminated. The mouse had to nose poke into any of the interfaces to receive the reward. The intertrial interval was 4 s and there was no time limit for the mouse to select one of the five interfaces. The training sessions for this phase were 30 min and the mouse advanced to 5CSRTT training after 2 consecutive sessions with >60 rewards claimed. The 5CSRTT training has four stages with a similar paradigm. At the start of the 30 min session, the mouse was given a free reward to initiate testing. One of the 5 interfaces on the testing wall was illuminated for a given stimulus duration (SD) based on the stage of training. The mouse had to nose poke into the correct hole within the limited hold (LH) period to receive the reward. The LH was based on the SD time. If the SD was 10 or 8 s, the LH was 10 s from the onset of the light, if the SD was 4 or 2 s, the LH was 5 s. There was a 4 s intertrial interval between correct responses. If the mouse made a premature nose poke, selected the incorrect interface, or made no selection (omission), the mouse was given a 4 s time out where the house light was extinguished, and no reward was delivered. If the mouse made a premature selection, it did not count towards the total number of trials. For the first stage of testing, the SD was set at 10 s and mice advanced to the next stage by making >30 correct choices with a mean correct response time of 5 s on two consecutive days. Subsequent stages reduced the SD time to 8, 4, and 2 s with the same criteria for advancement: >30 correct choices with a mean correct response time that was 50% of the SD (1.5 s for the 2 s SD) on two consecutive days. After 5CSRTT training, mice were tested for 3 days using a variable SD (vSD) paradigm followed by 3 days of testing with a variable ITI (vITI) paradigm. The SDs used for vSD testing were 0.2, 0.4, 0.8, and 1.6 s with a limited hold of 5 s and the ITI was held at 4 s. For vITI testing, the SD was held at 2 s, the LH was 5 s and the ITIs were 3, 4, 5, 6, and 7 s. The number of correct responses, incorrect responses, premature responses, omission to response, total number of entries into the reward hole, and nose pokes during the time out were recorded. Latency to commit both correct and incorrect responses was also recorded as was latency to collect the reward. Accuracy was calculated as a percentage using the following formula: (100 * correct responses) / (correct responses + incorrect responses). The percent of premature responses was calculated as: (100 x premature responses) / (premature responses + incorrect responses + correct responses). The percent of omissions was calculated as (100*omissions) / (total trials). The number of preservative responses, defined as a nose poke into the unlit hole that represented the correct selection on the previous trial, was recorded.

### Statistical analysis

The primary hypothesis of this study was that *Slc6a8* loss in the brain would be sufficient to disrupt 5CSRTT performance compared with control mice. Therefore, the first level of testing was the bKO mice compared with the combined control groups. The training, vITI, and vSD data were analyzed using mixed effects model ANOVAs using an autoregressive correlation structure. Group was the between subject factor, ITI or SD were repeated measures, and litter and day were blocking factors. Kenward-Rogers degrees of freedom were used and can result in different degrees of freedom for different datasets from the same mice. Day was used as a blocking factor since mixed models can only calculate one repeated measure and trial (ITI or SD) which was hypothesized to be a greater influence on performance than day. To determine if the FLOX and NES mice differed from controls, a second analysis was performed where each genotype was separated. For main effects of gene, the differences of LS Means were then calculated for each comparison. Cumulative Gaussian analysis was performed to determine the percent of mice that had completed training by a given day. The data were plotted using a frequency distribution plot and fit using a least squares regression and the curves were compared using the extra sum of squares F test. The F tables and differences of LS means are presented in tables to improve the presentation of the results. The differences of LS means for significant effects of gene are shown in the supplemental data.

## Results

*Body weight:* Mice were weighed prior to testing. There was a main effect of gene (F(3,29)=0.2916, P<0.01, not shown) for weight with NES mice weighing less than WT while there were no differences between other groups and the bKO mice did not differ from controls (Figure 1A). During the 5C-SRTT testing, there was a main effect of gene on the percent of weight reduction, however there were no individual difference between groups.

**Figure 1.**
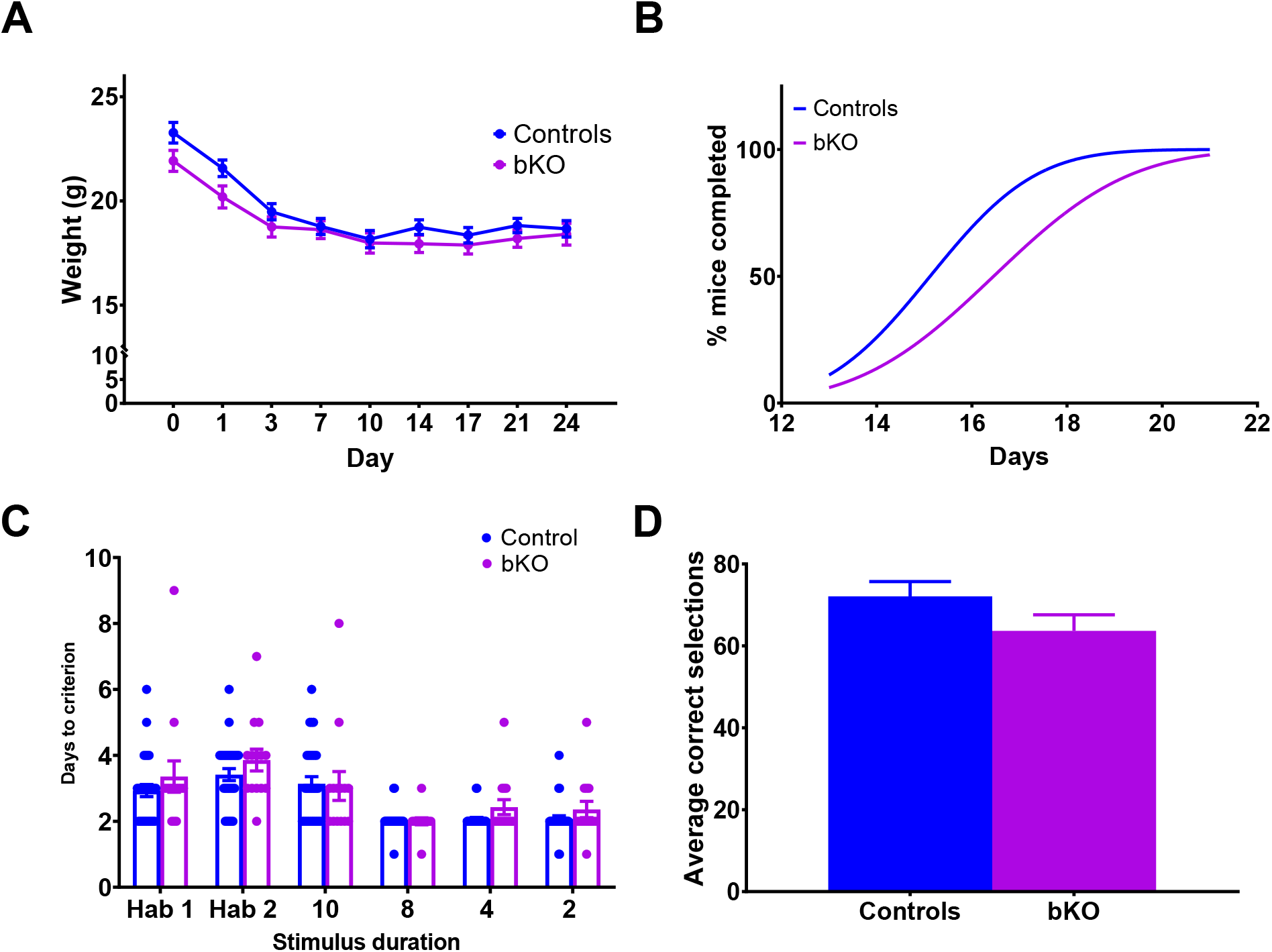
Body weight and training data. (A) The bKO mice do not show differences in body weight at start and tolerate weight restriction. (B) The cumulative distribution of days to complete the training phase of the 5CSRTT showing that bKO mice take longer to complete the training phase compared with control mice. (C) Time to complete the individual training phases in the 5CSRTT. (D) The number of correct responses during the HAB2 phase of testing, where all 5 lights were presented. N=29 controls; 14 bKO mice.

### Training

Table 1 shows the F values for all variables in this phase. *Combined controls:* It took more sessions for the mice in the bKO group to complete the training phase of the 5C-SRTT compared with controls (F(2,16)=36.64, P<0.001, Figure 1B). There were no differences in the number of days required to pass an individual phase of testing (Figure 1C). During the 2^nd^ phase of habituation testing, there was no difference in the number of correct responses between control and bKO mice and there were no individual differences between groups (Figure 1D). The bKO mice had more omissions across all SDs compared with the control group (Figure 2A). There was no difference in accuracy (Figure 2B) between groups. There was a main effect of gene and a gene x SD interaction for correct latency with bKO mice having longer latencies to make a correct response at 8 and 4 s SDs than the control group (Figure 2C). The bKO mice also took longer to obtain the reward compared with the control mice (Figure 2E). The bKO mice did not show differences in premature responses (Figure 2D), time out entries (Figure 2F), incorrect entries into the reward hole (Figure 2G), or perseverant responses (Figure 2H) compared with controls. There was a main effect of SD for all variables except reward entries and premature rate. *Split controls:* The same variables that showed main effects of gene when the controls were combined showed main effects of gene when controls were separated (Table 1). Analysis of the differences of LSMeans showed no difference between control groups at the level of gene for any variable analyzed. However, there were gene X SD interactions for correct latency, perseverative entries, and reward hole entries (Supplementary Figure 1). During the 8 s SD phase, the NES, FLOX, and bKO mice had longer latencies to make a correct selection compared with WT mice. The bKO mice also had longer correct latencies than NES and FLOX mice while the NES and FLOX mice did not differ. The FLOX and bKO mice had more head entries than WT and NES mice during the 2 s SD phase. During the 8 s SD phase, the WT mice had fewer perseverative entries than all other groups, which did not differ from one another.

**Table 1.** F values for variables during the 5 choice training.

|  |  | *Combined Controls* |  |  |  | *Split controls* |  |  |  |
| --- | --- | --- | --- | --- | --- | --- | --- | --- | --- |
| **Variable** | **Factor** | **DF** | **F** | **P** | **Sig.** | **DF** | **F** | **P** | **Sig.** |
| **%Omissions** | *Gene* | 1, 44.8 | 11.39 | 0.0015 | ** | 3, 43 | 5.04 | 0.0044 | ** |
|  | *SD* | 3, 114 | 22.26 | <0.0001 | *** | 3, 108 | 27.66 | <0.0001 | *** |
|  | *Gene\*SD* | 3, 114 | 1.1 | 0.3520 |  | 9, 113 | 0.86 | 0.5634 |  |
| **% Correct** | *Gene* | 1, 49.8 | 8.57 | 0.0051 | ** | 3, 46.2 | 5.05 | 0.0041 | ** |
|  | *SD* | 3, 121 | 58.14 | <0.0001 | *** | 3, 113 | 62.16 | <0.0001 | *** |
|  | *Gene\*SD* | 3, 121 | 0.32 | 0.8086 |  | 9, 117 | 1.09 | 0.3720 |  |
| **Accuracy** | *Gene* | 1, 47.7 | 0.67 | 0.4183 |  | 3, 44.3 | 0.89 | 0.4533 |  |
|  | *SD* | 3, 119 | 42.40 | <0.0001 | *** | 3, 113 | 39.26 | <0.0001 | *** |
|  | *Gene\*SD* | 3, 119 | 0.76 | 0.5209 |  | 9, 115 | 0.82 | 0.6028 |  |
| **Premature Rate** | *Gene* | 1, 54.9 | 2.06 | 0.1570 |  | 3, 51.8 | 1.03 | 0.3850 |  |
|  | *SD* | 3, 121 | 0.32 | 0.8112 |  | 3, 115 | 0.66 | 0.5803 |  |
|  | *Gene\*SD* | 3, 121 | 1.24 | 0.2980 |  | 9, 120 | 0.79 | 0.6243 |  |
| **Correct Latency** | *Gene* | 1, 45.6 | 9.18 | 0.0040 | ** | 3, 43.1 | 3.89 | 0.0152 | * |
|  | *SD* | 3, 114 | 290.7 | <0.0001 | *** | 3, 108 | 311.20 | <0.0001 | *** |
|  | *Gene\*SD* | 3, 114 | 4.26 | 0.0068 | ** | 9, 113 | 1.56 | 0.1354 |  |
| **Latency to Reward** | *Gene* | 1, 41.1 | 8.07 | 0.0070 | ** | 3, 39 | 3.81 | 0.0174 | * |
|  | *SD* | 3, 114 | 34.94 | <0.0001 | *** | 3, 108 | 36.64 | <0.0001 | *** |
|  | *Gene\*SD* | 3, 114 | 0.52 | 0.6697 |  | 9, 111 | 0.57 | 0.8515 |  |
| **Time out entries** | *Gene* | 1, 48.1 | 0.13 | 0.6305 |  | 3, 45 | 1.10 | 0.3583 |  |
|  | *SD* | 3, 118 | 35.74 | <0.0001 | *** | 3, 112 | 47.50 | <0.0001 | *** |
|  | *Gene\*SD* | 3, 118 | 1.56 | 0.2027 |  | 9, 115 | 1.21 | 0.2944 |  |
| **Reward hole Entries** | *Gene* | 1, 41.1 | 0.23 | 0.5284 |  | 3, 39.8 | 2.13 | 0.1119 |  |
|  | *SD* | 3, 118 | 0.98 | 0.4059 |  | 3, 113 | 1.61 | 0.1907 |  |
|  | *Gene\*SD* | 3, 118 | 0.19 | 0.9061 |  | 9, 113 | 3.16 | 0.0019 | ** |
| **% Incorrect** | *Gene* | 1, 50.9 | 0.06 | 0.8119 |  | 3, 47.5 | 0.35 | 0.7880 |  |
|  | *SD* | 3, 119 | 17.59 | <0.0001 | *** | 3, 113 | 14.04 | <0.0001 | *** |
|  | *Gene\*SD* | 3, 119 | 0.97 | 0.4112 |  | 9, 117 | 0.92 | 0.5148 |  |
| **Perseverant Responses** | *Gene* | 1, 59.3 | 1.68 | 0.2001 |  | 3, 54 | 2.39 | 0.0790 |  |
|  | *SD* | 3, 126 | 3.20 | 0.0256 | * | 3, 118 | 4.00 | 0.0094 | ** |
|  | *Gene\*SD* | 3, 126 | 0.47 | 0.7043 |  | 9, 123 | 2.37 | 0.0167 | * |

**Figure 2.**
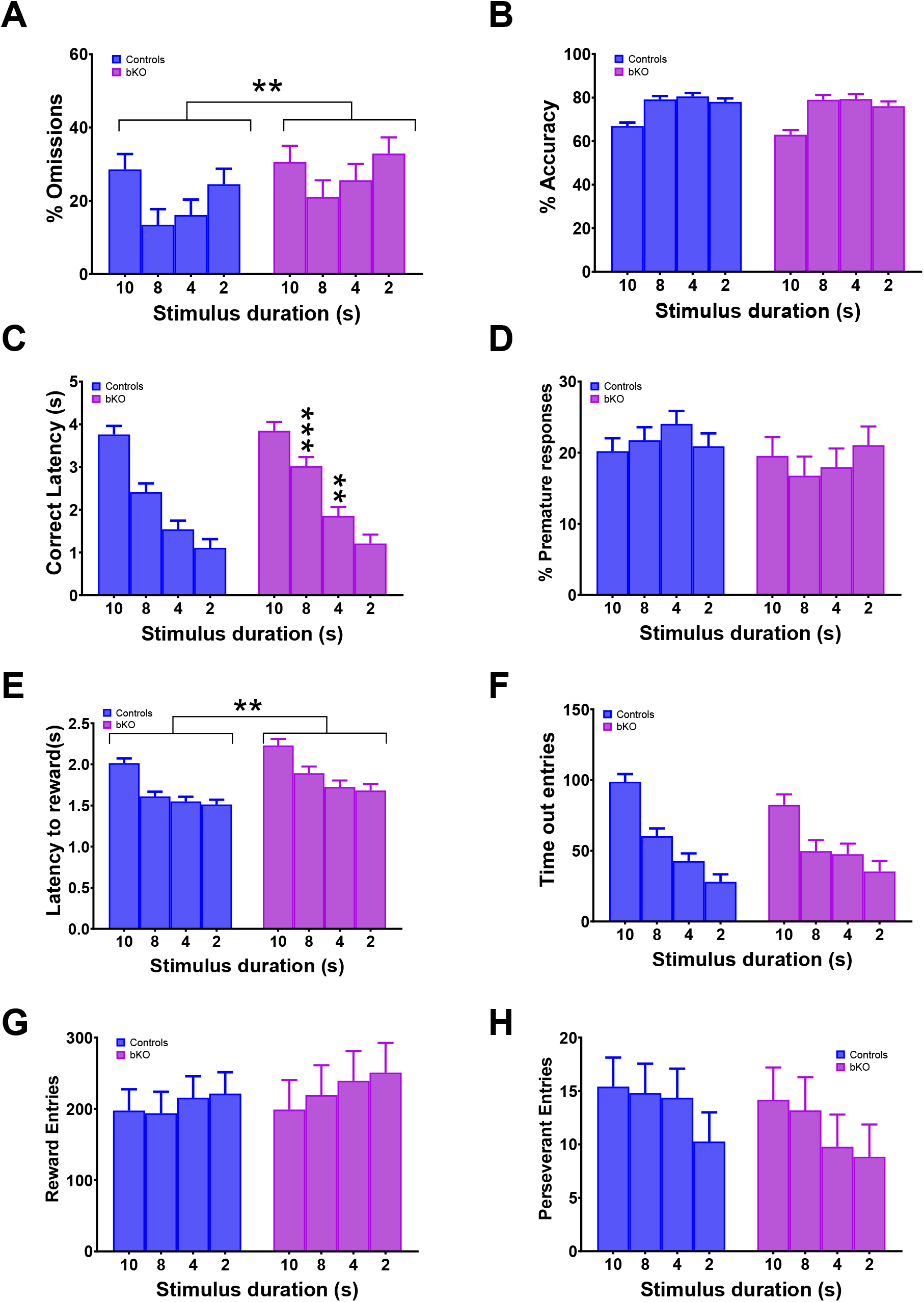
5CSRTT training phase. The bKO mice have an increase in omissions (A) and longer latencies to reward (E) compared with controls. There was an increase in the latency to make the correct choice (C) during the phases of testing with an 8 or 4 s SD. No changes were observed in any other measures. **P>0.01 bKO compared with controls; ***P>0.001 bKO vs controls. Brackets indicate a main effect of gene. N=29 controls; 14 bKO mice.

### Testing-Variable SD

Table 2 shows the F table for all variables. *Combined controls:* The bKO mice had more omissions compared with controls (Figure 3A). They also showed a reduction in accuracy (Figure 3B) and had slower latencies to make the correct choice (Figure 3C) and obtain the reward (Figure 3E) than controls. The bKO mice committed more nose pokes during the time out periods compared with controls (Figure 3F). No changes were observed in premature or perseverant responses in the bKO mice compared with controls. There were no interactions of gene and cue duration for any variable during this phase of testing. *Split controls:* The only variable to show a main effect of gene in the split control ANOVA but not the combined control ANOVA was the rate of premature entries. For omissions, accuracy, latency to reward, and correct latency, there were no differences between control groups. Analysis of differences of LSMeans showed that NES and bKO mice had a higher rate of premature entries compared with FLOX mice. For time out entries, the NES, FLOX, and bKO mice had more entries than WT mice and did not differ from one another. The FLOX mice had more reward entries than the WT and NES mice, but FLOX and bKO were similar. The bKO mice had more reward entries than the WT mice, but not the NES mice.

**Table 2.** F values for variables in vSD testing

| Variable | Factor | *Combined Controls* |  |  |  | *Split Controls* |  |  |  |
| --- | --- | --- | --- | --- | --- | --- | --- | --- | --- |
|  |  | DF | F | P | Sig. | DF | F | P | Sig. |
| % Omissions | *Gene* | 1, 131 | 16.51 | <0.0001 | *** | 3, 129 | 6.36 | 0.0005 | *** |
|  | *Cue* | 3, 346 | 38.33 | <0.0001 | *** | 3, 340 | 38.61 | <0.0001 | *** |
|  | *Gene\*Cue* | 3, 346 | 0.20 | 0.8960 |  | 9, 348 | 1.12 | 0.3478 |  |
| % Correct | *Gene* | 1, 143 | 20.04 | <0.0001 | *** | 3, 140 | 6.94 | 0.0002 | *** |
|  | *Cue* | 3, 349 | 171.53 | <0.0001 | *** | 3, 343 | 187.64 | <0.0001 | *** |
|  | *Gene\*Cue* | 3, 349 | 0.34 | 0.7938 |  | 9, 356 | 0.97 | 0.4623 |  |
| Accuracy | *Gene* | 1, 138 | 7.97 | 0.0054 | ** | 3, 155 | 4.02 | 0.0087 | ** |
|  | *Cue* | 3, 346 | 149.50 | <0.0001 | *** | 3, 341 | 157.44 | <0.0001 | *** |
|  | *Gene\*Cue* | 3, 346 | 0.51 | 0.6777 |  | 9, 360 | 0.44 | 0.9129 |  |
| Premature Rate | *Gene* | 1, 126 | 0.15 | .06978 |  | 3, 125 | 3.45 | 0.0186 | * |
|  | *Cue* | 3, 341 | 0.85 | 0.4655 |  | 3, 335 | 1.44 | 0.2321 |  |
|  | *Gene\*Cue* | 3, 341 | 1.17 | 0.3217 |  | 9, 344 | 0.60 | 0.7928 |  |
| Correct latency | *Gene* | 1, 172 | 5.65 | 0.0185 | *** | 3, 164 | 1.98 | 0.1195 |  |
|  | *Cue* | 3, 333 | 2.34 | 0.0731 | *** | 3, 327 | 3.49 | 0.0161 | * |
|  | *Gene\*Cue* | 3, 333 | 0.87 | 0.4555 |  | 9, 355 | 0.57 | 0.8238 |  |
| Reward Latency | *Gene* | 1, 122 | 15.20 | 0.0002 | *** | 3, 119 | 6.31 | 0.0005 | *** |
|  | *Cue* | 3, 340 | 0.29 | 0.8351 | * | 3, 335 | 0.19 | 0.9063 |  |
|  | *Gene\*Cue* | 3, 340 | 1.02 | 0.3843 |  | 9, 341 | 0.55 | 0.8389 |  |
| Time out entries | *Gene* | 1, 121 | 6.31 | 0.0133 | ** | 3, 120 | 4.61 | 0.0043 | ** |
|  | *Cue* | 3, 337 | 7.87 | <0.0001 | *** | 3, 331 | 9.46 | <0.0001 | *** |
|  | *Gene\*Cue* | 3, 337 | 0.70 | 0.5520 |  | 9, 340 | 0.84 | 0.5789 |  |
| Receptacle entries | *Gene* | 1, 117 | 0.22 | 0.6363 |  | 3, 117 | 3.98 | 0.0097 | ** |
|  | *Cue* | 3, 346 | 2.68 | 0.0471 | ** | 3, 332 | 3.86 | 0.0098 | ** |
|  | *Gene\*Cue* | 3, 346 | 1.42 | 0.2381 |  | 9, 334 | 2.37 | 0.0131 | ** |
| % Incorrect | *Gene* | 1, 134 | 1.13 | 0.2899 |  | 3, 132 | 1.37 | 0.2539 |  |
|  | *Cue* | 3, 339 | 47.82 | <0.0001 | *** | 3, 334 | 58.25 | <0.0001 | *** |
|  | *Gene\*Cue* | 3, 339 | 0.96 | 0.4111 |  | 9, 348 | 0.79 | 0.6298 |  |
| Perseverant | *Gene* | 1, 146 | 0.14 | 0.7050 |  | 3, 143 | 0.65 | 0.5851 |  |
|  | *Cue* | 3, 333 | 6.40 | 0.0003 | *** | 3, 328 | 5.41 | 0.0012 | ** |
|  | *Gene\*Cue* | 3, 333 | 0.26 | 0.8561 |  | 9, 349 | 0.89 | 0.5347 |  |

**Figure 3.**
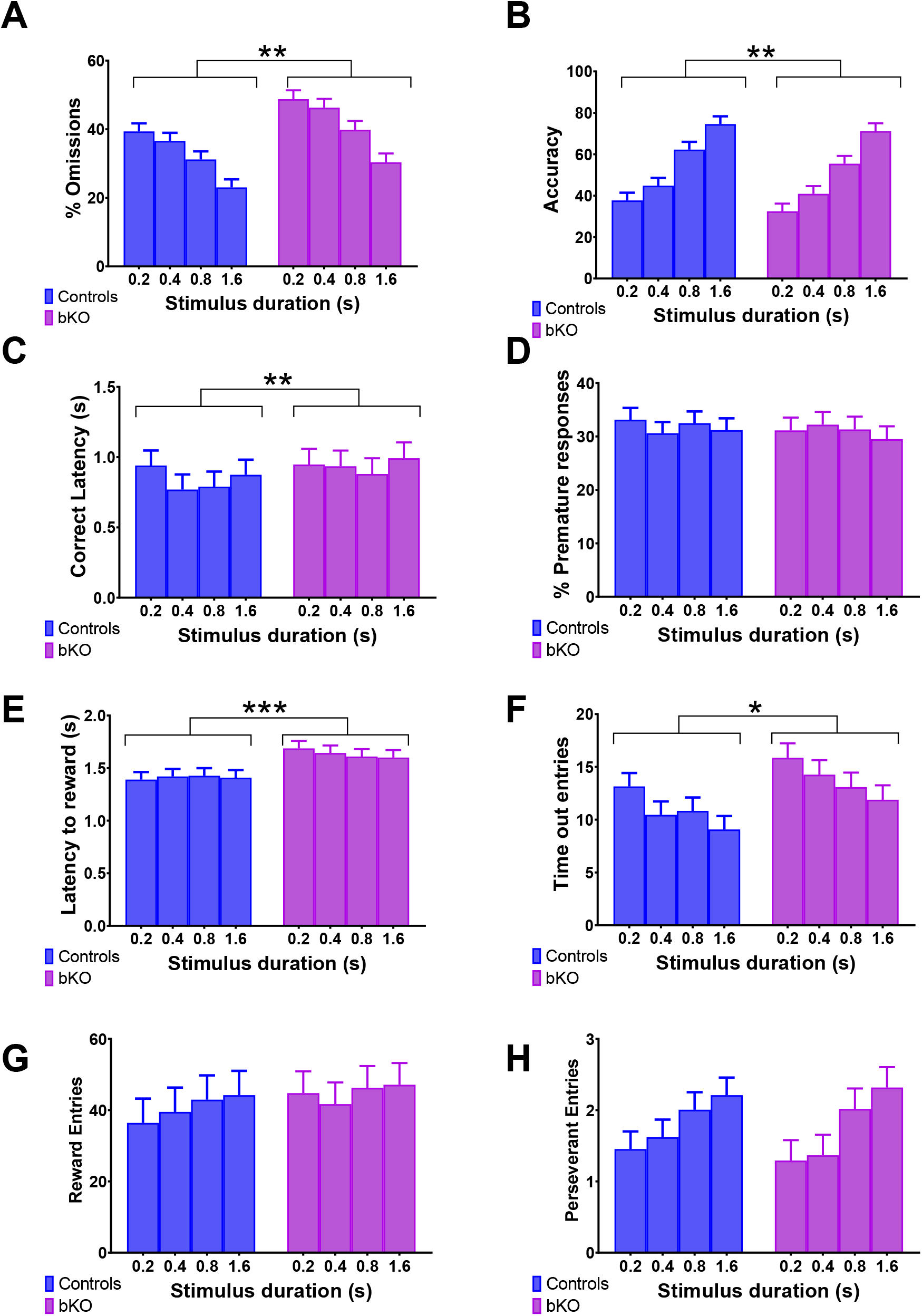
Variable stimulus duration testing. The bKO mice show increases in the percent of omissions (A), reduced accuracy (B), and an increase in correct latency (C) compared with control mice. The bKO mice also show increases in latency to obtain the reward (E) and the number of nose pokes during the time out period (F). There were no changes in premature responses (D), reward hole entries (G), perseverant entries (H). *P>0.05; **P>0.01; ***P>0.001 bKO vs controls. Brackets indicate a main effect of gene. N=29 controls; 14 bKO mice.

### Testing-Variable ITI

The F values for each ANOVA are shown in Table 3. *Combined controls:* The bKO mice committed more omissions (Figure 4A) and had a lower accuracy (Figure 4B) compared with controls. The bKO mice had longer latencies to both make a correct choice (Figure 4C) and claim the reward (Figure 4E) following a correct choice than controls. Time out entries (Figure 4F), and erroneous reward hole entries (Figure 4G) were increased in the bKO mice compared with controls while the number of perseverant responses (Figure 4H) was reduced. The premature rate was unchanged between groups. As with the vSD phase, there were no interactions of gene and ITI duration during this phase. There were main effects of ITI for every variable except perseverant responses. *Split controls:* Similar main effects of gene were seen between the split and combined control analysis. Differences between control groups were observed in reward latency and time out entries. Both the bKO and NES mice had longer reward latencies compared with WT and FLOX mice. The FLOX and bKO mice had more time out entries compared with the WT and NES mice, which did not differ.

**Table 3.** F values for variable ITI testing.

| Variable | Factor | *Combined Controls* |  |  |  | *Split Controls* |  |  |  |
| --- | --- | --- | --- | --- | --- | --- | --- | --- | --- |
|  |  | DF | F | P | Sig. | DF | F | P | Sig |
| % Omissions | *Gene* | 1, 136 | 16.81 | <0.0001 | *** | 3, 134 | 5.85 | 0.0009 | *** |
|  | *ITI* | 4, 478 | 18.59 | <0.0001 | *** | 4, 470 | 15.77 | <0.0001 | *** |
|  | *Gene\*ITI* | 4, 478 | 2.08 | 0.0829 |  | 12, 486 | 1.17 | 0.3040 |  |
| % Correct | *Gene* | 1, 147 | 19.51 | <0.0001 | *** | 3, 144 | 6.57 | 0.0003 | *** |
|  | *ITI* | 4, 476 | 5.22 | 0.0004 | *** | 4, 468 | 5.11 | 0.0005 | *** |
|  | *Gene\*ITI* | 4, 476 | 1.16 | 0.3256 |  | 12, 490 | 0.81 | 0.6396 |  |
| Accuracy | *Gene* | 1, 168 | 4.80 | 0.0298 | * | 3, 161 | 1.59 | 0.1930 |  |
|  | *ITI* | 4, 484 | 10.33 | <0.0001 | *** | 4, 476 | 10.85 | <0.0001 | *** |
|  | *Gene\*ITI* | 4, 484 | 0.52 | 0.7227 |  | 12, 502 | 0.77 | 0.6849 |  |
| Premature Rate | *Gene* | 1, 130 | 2.16 | 0.1437 |  | 3, 124 | 0.72 | 0.5425 |  |
|  | *ITI* | 4, 479 | 82.09 | <0.0001 | *** | 4, 471 | 88.38 | <0.0001 | *** |
|  | *Gene\*ITI* | 4, 479 | 0.99 | 0.4115 |  | 12, 482 | 0.53 | 0.7893 |  |
| Correct latency | *Gene* | 1, 166 | 25.49 | <0.0001 | *** | 3, 164 | 9.66 | <0.0001 | *** |
|  | *ITI* | 4, 464 | 33.27 | <0.0001 | *** | 4, 456 | 35.16 | <0.0001 | *** |
|  | *Gene\*ITI* | 4, 464 | 0.59 | 0.6719 |  | 12, 492 | 1.01 | 0.4408 |  |
| Reward Latency | *Gene* | 1, 135 | 5.86 | 0.0168 | * | 3, 132 | 10.67 | <0.0001 | *** |
|  | *ITI* | 4, 491 | 2.41 | 0.0487 | * | 4, 481 | 2.45 | 0.0452 | * |
|  | *Gene\*ITI* | 4, 491 | 0.56 | 0.6926 |  | 12, 488 | 1.24 | 0.2509 |  |
| Time out entries | *Gene* | 1, 121 | 7.52 | 0.0070 | ** | 3, 118 | 4.05 | 0.0088 | ** |
|  | *ITI* | 4, 466 | 18.98 | <0.0001 | *** | 4, 458 | 15.20 | <0.0001 | *** |
|  | *Gene\*ITI* | 4, 466 | 1.34 | 0.2526 |  | 12, 472 | 1.08 | 0.3728 |  |
| Receptacle entries | *Gene* | 1, 127 | 3.94 | 0.0492 | * | 3, 119 | 5.46 | 0.0015 | ** |
|  | *ITI* | 4, 482 | 3.08 | 0.0159 | * | 4, 474 | 2.88 | 0.0224 | ** |
|  | *Gene\*ITI* | 4, 482 | 0.63 | 0.6393 |  | 12, 476 | 0.86 | 0.5845 |  |
| % Incorrect | *Gene* | 1, 173 | 0.30 | 0.5877 |  | 3, 170 | 0.19 | 0.9023 |  |
|  | *ITI* | 4, 487 | 14.21 | <0.0001 | *** | 4, 480 | 14.64 | <0.0001 | *** |
|  | *Gene\*ITI* | 4, 487 | 0.64 | 0.6351 |  | 12, 505 | 0.80 | 0.6471 |  |
| Perseverant | *Gene* | 1, 141 | 4.78 | 0.0304 | * | 3, 136 | 2.05 | 0.1095 |  |
|  | *ITI* | 4, 491 | 0.43 | 0.7854 |  | 4, 483 | 1.52 | 0.1952 |  |
|  | *Gene\*ITI* | 4, 491 | 1.82 | 0.1235 |  | 12, 492 | 1.35 | 0.1866 |  |

**Figure 4.**
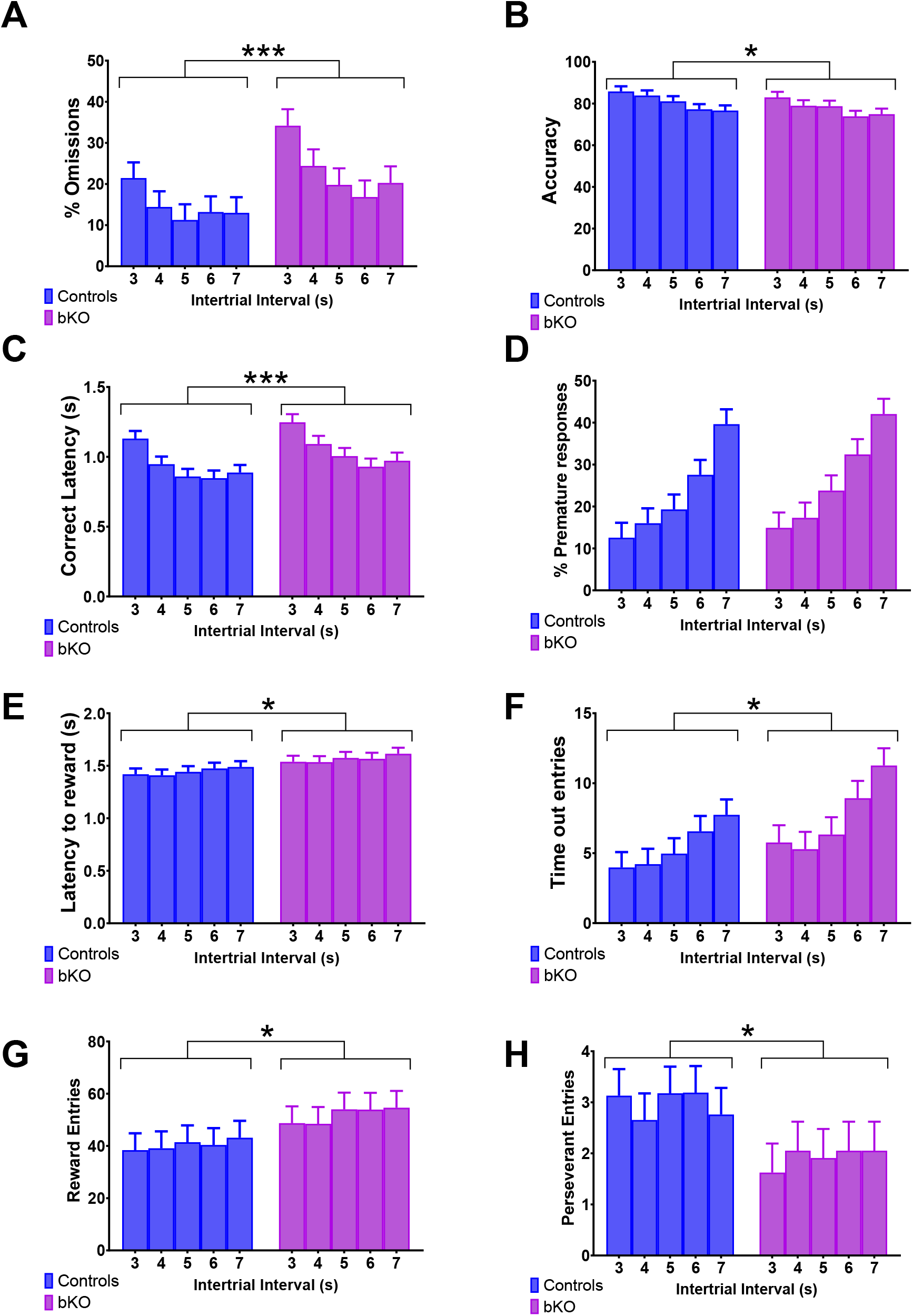
Variable ITI testing. The bKO mice show increases in the percent of omissions (A), reduced accuracy (B), and an increase in correct latency (C) compared with control mice. There were no differences in premature responses (D). The latency to claim the reward (E), the number of time out entries (F), and the number of reward hole entries (G) were increased in bKO mice while the number of perseverant entries (H) were lower. *P>0.05; **P>0.01; ***P>0.001 bKO vs controls. Brackets indicate a main effect of gene. N=29 controls; 14 bKO mice.

## Discussion

We have shown that mice lacking brain Cr have deficits in spatial learning, object recognition memory, and conditioned memories ^3,17-19,28^. While these findings suggest that *Slc6a8*^*-/y*^ mice are a high-fidelity model of CTD, there is a need to expand our understanding of the nature of the CTD deficit and test the mice in a more translationally relevant cognitive test. In this study, we show for the first time that mice lacking brain Cr display an inattentive-like phenotype in the 5C-SRTT. The bKO mice show reductions in accuracy and processing speed along with an increase in the percent of omitted trials across both test phases of the 5C-SRTT.

The bKO mice were used in this study because the small stature of the *Slc6a8*^*-/y*^ mice led to mortality during food deprivation and concerns that the smaller stature of the *Slc6a8*^*-/y*^ mice could confound any changes observed. We have shown that the bKO mice recapitulate many of the learning deficits seen in the *Slc6a8*^*-/y*^ mice while not disturbing the physical characteristics of the mice ^18^. In agreement with this study, there were no weight differences between bKO and control mice at the start of testing. The bKO mice were able to complete the training phases of the task; however, they took longer to complete the training phase of the 5C-SRTT than control mice. The increase in days required to complete the task do not appear to be due to a single phase of training, rather an accumulation of extra days across all trials. The bKO mice show an increase in omissions during the testing phase along with longer correct latencies in the middle phases, suggesting that they have a pervasive inattentive phenotype even at longer SDs.

Shorter SD’s are designed to test attention using more demanding parameters ^24,27^. As can be seen in Figure 3, omissions increased, and accuracy was reduced for all mice as the SD was shortened. The bKO mice showed an increase in omissions, reduced accuracy, and longer correct latencies that was not dependent on the SD. These findings are consistent with an inattentive phenotype. This is reinforced during the vITI phases, where the bKO mice have the same changes in omissions, accuracy, and correct latency. The vITI phase is used to assess impulsive-like behaviors. As can be seen, the number of premature responses increase as the ITI gets longer. There were no differences between bKO and control mice, suggesting that the bKO mice do not have an impulsive like phenotype. This interpretation could be confounded by the increase in the number of nose pokes during the time out, though this finding could be a function of the bKO mice receiving more time outs than control mice. The reduction in perseverative entries, nose poking into the previous target hole, would suggest that the bKO mice have lower compulsive behaviors as well, suggesting that the primary phenotype of these mice is inattention. These findings are in agreement with limited data from CTD patients, with many (55%) showing ADD/ADHD while impulsive (27%) or compulsive (8%) behaviors were not as common ^29^.

While this is the first study to evaluate attention and impulsivity in Cr-deficient rodent models, other aspects of executive function have been evaluated. Reductions in spontaneous alternation, a measurement of working memory using the Y-maze, have been consistently seen in mice with exons 5-7 removed from the *Slc6a8* gene ^20,30,31^. The reductions in spontaneous alternations have also been seen in a rat model of CTD ^21^. Spontaneous alternation has not been reported using our mouse model, but we have recently discovered that *Slc6a8*^*-/y*^ mice do not have working memory deficits in a swimming version of the radial arm maze (Sugimoto et al, *in preparation*). To our knowledge, there has not been a comprehensive evaluation of executive function in CTD patients, so it is difficult to fully determine the translational relevance of these findings.

While the primary hypothesis of this study was to evaluate the effect of *Slc6a8* deletion in the brain of mice, it is also important to ensure that any deficits observed are due to *Slc6a8* deletion instead of the contribution of the *Nestin-*Cre expressing transgenic mouse. Deficits in conditioned fear have been observed in *Nestin-*Cre mice, with no differences seen in other behaviors examined ^32^. The *Nestin-*Cre mice did not differ from WT mice in any of the primary measurements-accuracy, omissions, correct latency, and premature responses during any phase. During the vSD phase, only difference between WT and *Nestin-*Cre mice was time out entries, with the *Nestin-*Cre mice having more entries. Interestingly, the FLOX mice also had more entries than the WT, but the bKO did not differ from either. While it also important to determine if the floxed *Slc6a8* mice contribute to the deficit, any changes seen in the FLOX mice are likely due to mild changes in *Slc6a8* function. During the vITI testing, the *Nestin-Cre* mice had longer reward latencies than WT mice. Together, while there were some changes between the control mice, it appears that they did not contribute to the changes seen in the bKO mice.

In conclusion, we have shown that mice that lack the *Slc6a8* gene in the brain show an inattentive-like phenotype compared with controls. The results of this study provide additional, translationally relevant data to advance the understanding of Cr in the brain as well as provide another avenue in which to test potential therapies.

## Author contributions

MRS designed the experiments, participated in the execution of the experiments, analyzed the data, and wrote the manuscript. RL participated in the execution of the experiments. MKP participated in the design and execution of the experiments and edited the manuscript.

## Ethics statement

All experiments conducted on mice were approved by the Institutional Animal Care and Use Committee of Cincinnati Children’s Research Foundation. IACUC protocol number 2020-0009.

**The authors declare no conflicts of interest**.

## Data availability

All data will be made available upon request to MRS.

