## Supplemental tables and figures for "Creatine transporter (Slc6a8) knockout mice show an inattentive-like phenotype in the 5-choice serial reaction time test"

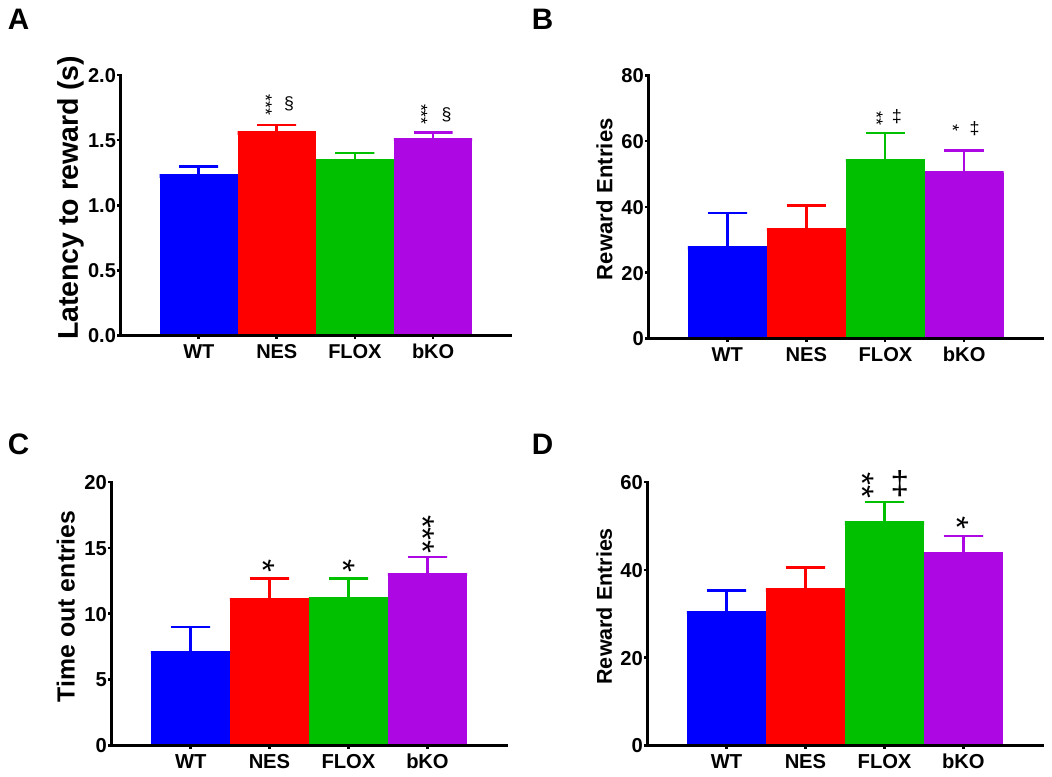

**Supplemental Figure 1. Significant differences between control groups.** (A) Latency to reward during vITI testing was increased for the NES and bKO mice compared with FLOX and WT. (B) Reward entries were increased for FLOX and bKO mice during vITI testing. (C) Time out entries were increased for all groups compared with WT during vSD testing. (D) FLOX an bKO mice showed more reward entries during the vSD testing. *,**, *** p<0.05, 0.01, 0.001 vs WT, respectively; ‡ p<0.05 vs NES; § p<0.05 vs FLOX.

| **Variable** | **Comparison** | **DF** | **t** | **P** | **Sig.** |
| --- | --- | --- | --- | --- | --- |
| % Omission | WT-NES | 39.2 | -1.15 | 0.258 |  |
|  | WT-FLOX | 39.3 | -1.23 | 0.2257 |  |
|  | WT-bKO | 38.2 | -3.35 | 0.0018 | ** |
|  | NES-FLOX | 39.8 | -0.2 | 0.8419 |  |
|  | NES-bKO | 38.8 | -2.34 | 0.0245 | * |
|  | FLOX-bKO | 39 | -1.87 | 0.0692 |  |
| % Correct | WT-NES | 40.1 | 1.11 | 0.2722 |  |
|  | WT-FLOX | 40.2 | 2.28 | 0.0282 | ** |
|  | WT-bKO | 39.2 | 3.58 | 0.0009 | *** |
|  | NES-FLOX | 40.7 | 1.34 | 0.1862 |  |
|  | NES-bKO | 39.7 | 2.62 | 0.0123 | * |
|  | FLOX-bKO | 39.9 | 0.94 | 0.3547 |  |
| Latency to Reward | WT-NES | 39 | -1.25 | 0.2201 |  |
|  | WT-FLOX | 38.8 | 0.42 | 0.6766 |  |
|  | WT-bKO | 38.9 | -2.6 | 0.0132 | * |
|  | NES-FLOX | 39 | 1.65 | 0.1067 |  |
|  | NES-bKO | 39.2 | -1.42 | 0.163 | * |
|  | FLOX-bKO | 38.9 | -2.96 | 0.0052 | ** |

Supplemental table 1. Differences of LS Means for main effect of gene during training trials.

| **Variable** | **Comparison** | **DF** | **t** | **P** | **Sig.** |
| --- | --- | --- | --- | --- | --- |
| % Omission | WT-NES | 130 | 1.41 | 0.1613 |  |
|  | WT-FLOX | 130 | 0.32 | 0.7492 |  |
|  | WT-bKO | 130 | -2.3 | 0.0229 | * |
|  | NES-FLOX | 129 | -1.21 | 0.2277 |  |
|  | NES-bKO | 129 | -4.09 | <.0001 | *** |
|  | FLOX-bKO | 129 | -2.99 | 0.0033 | ** |
| % Correct | WT-NES | 141 | -0.48 | 0.6306 |  |
|  | WT-FLOX | 141 | 0.25 | 0.8008 |  |
|  | WT-bKO | 137 | 3.12 | 0.0022 | ** |
|  | NES-FLOX | 140 | 0.8 | 0.4252 |  |
|  | NES-bKO | 140 | 3.96 | 0.0001 | *** |
|  | FLOX-bKO | 140 | 3.31 | 0.0012 | ** |
| Accuracy | WT-NES | 155 | 1.53 | 0.1281 |  |
|  | WT-FLOX | 155 | 1.47 | 0.1438 |  |
|  | WT-bKO | 155 | 3.4 | 0.0009 | *** |
|  | NES-FLOX | 155 | -0.14 | 0.886 |  |
|  | NES-bKO | 155 | 1.69 | 0.0932 |  |
|  | FLOX-bKO | 155 | 1.97 | 0.0508 |  |
| Premature Rate | WT-NES | 125 | -1.96 | 0.0519 | * |
|  | WT-FLOX | 125 | 0.98 | 0.3309 |  |
|  | WT-bKO | 125 | -0.01 | 0.9933 |  |
|  | NES-FLOX | 124 | 3.17 | 0.0019 | ** |
|  | NES-bKO | 124 | 2.27 | 0.0247 | ** |
|  | FLOX-bKO | 124 | -1.15 | 0.2504 |  |
| Reward Latency | WT-NES | 120 | -1.7 | 0.0918 |  |
|  | WT-FLOX | 120 | -1.45 | 0.1509 |  |
|  | WT-bKO | 117 | -4.06 | <.0001 | *** |
|  | NES-FLOX | 119 | 0.37 | 0.7124 |  |
|  | NES-bKO | 119 | -2.41 | 0.0176 | * |
|  | FLOX-bKO | 119 | -2.98 | 0.0035 | * |
| Time out entries | WT-NES | 120 | -2.33 | 0.0215 | * |
|  | WT-FLOX | 120 | -2.51 | 0.0135 | * |
|  | WT-bKO | 117 | -3.72 | 0.0003 | *** |
|  | NES-FLOX | 120 | -0.04 | 0.965 |  |
|  | NES-bKO | 120 | -1.3 | 0.1976 |  |
|  | FLOX-bKO | 120 | -1.33 | 0.1857 |  |
| Receptacle entries | WT-NES | 117 | -0.77 | 0.4452 |  |
|  | WT-FLOX | 117 | -3.15 | 0.002 | ** |
|  | WT-bKO | 117 | -2.26 | 0.0255 | * |
|  | NES-FLOX | 117 | -2.37 | 0.0196 | * |
|  | NES-bKO | 117 | -1.41 | 0.1621 |  |
|  | FLOX-bKO | 117 | 1.21 | 0.2298 |  |

**Supplemental Table 2. Differences of LS Means for main effect of gene during varSD testing.**

| **Variable** | **Comparison** | **DF** | **t** | **P** | **Sig.** |
| --- | --- | --- | --- | --- | --- |
| % Omission | WT-NES | 134 | -0.94 | 0.3469 |  |
|  | WT-FLOX | 134 | -0.39 | 0.695 |  |
|  | WT-bKO | 134 | -3.43 | 0.0008 | *** |
|  | NES-FLOX | 133 | 0.62 | 0.5342 |  |
|  | NES-bKO | 133 | -2.73 | 0.0071 | ** |
|  | FLOX-bKO | 133 | -3.4 | 0.0009 | *** |
| % Correct | WT-NES | 145 | 0.65 | 0.5161 |  |
|  | WT-FLOX | 144 | 0.08 | 0.9364 |  |
|  | WT-bKO | 145 | 3.38 | 0.0009 | *** |
|  | NES-FLOX | 143 | -0.64 | 0.5222 |  |
|  | NES-bKO | 143 | 3.06 | 0.0026 | ** |
|  | FLOX-bKO | 143 | 3.74 | 0.0003 | *** |
| Correct Latency | WT-NES | 165 | 1.72 | 0.088 |  |
|  | WT-FLOX | 165 | 0.65 | 0.5145 |  |
|  | WT-bKO | 165 | -2.62 | 0.0097 | ** |
|  | NES-FLOX | 164 | -1.21 | 0.2285 |  |
|  | NES-bKO | 163 | -5.06 | <.0001 | *** |
|  | FLOX-bKO | 164 | -3.73 | 0.0003 | *** |
| Reward Latency | WT-NES | 132 | -4.83 | <.0001 | *** |
|  | WT-FLOX | 132 | -1.73 | 0.0861 |  |
|  | WT-bKO | 132 | -4.51 | <.0001 | *** |
|  | NES-FLOX | 131 | 3.51 | 0.0006 | *** |
|  | NES-bKO | 131 | 0.73 | 0.4661 |  |
|  | FLOX-bKO | 131 | -3.07 | 0.0026 | ** |
| Time out entries | WT-NES | 118 | -1.76 | 0.0803 |  |
|  | WT-FLOX | 118 | -1.93 | 0.0554 |  |
|  | WT-bKO | 118 | -3.47 | 0.0007 | *** |
|  | NES-FLOX | 118 | -0.18 | 0.8568 |  |
|  | NES-bKO | 118 | -1.69 | 0.0938 |  |
|  | FLOX-bKO | 118 | -1.49 | 0.1379 |  |
| Receptacle entries | WT-NES | 108 | -0.65 | 0.5172 |  |
|  | WT-FLOX | 123 | -3.16 | 0.002 | ** |
|  | WT-bKO | 106 | -2.79 | 0.0063 | ** |
|  | NES-FLOX | 125 | -2.72 | 0.0074 | ** |
|  | NES-bKO | 124 | -2.51 | 0.0134 | * |
|  | FLOX-bKO | 125 | 0.53 | 0.5975 |  |

**Supplemental Table 3. Differences of LS Means for main effect of gene during varITI testing.**
